# jsdmstan: An R package for fitting joint species distribution models in Stan

**DOI:** 10.1101/2025.11.10.687559

**Authors:** Fiona Seaton

## Abstract

Joint species distribution models (JSDMs) have become an increasingly utilised tool for modelling and predicting change within communities of species across environmental gradients. Here we present a new R package for the fitting of JSDMs using the Bayesian probabilistic programming language Stan. The jsdmstan package can model species responses to environmental covariates and also species interactions through either full specification of the species covariance matrix or through a latent variable formulation. It also provides tools for simulating joint species distribution data according to those models. It supports specification of prior distributions of all parameters in the model, as well as access to the full suite of Stan diagnostics. The ability of this package to fit these models is demonstrated upon two real data sets, one on tree species in a survey of broadleaved woodlands and another on dune spider populations. The models are able to successfully recover population characteristics such as richness and show ecologically interesting results regarding residual species correlations and responses to environmental factors. The jsdmstan package provides a user-friendly interface for fitting JSDMs, with tools available to better understand both the consequences of the model assumptions and how well the model is performing.

## Introduction

Joint species distribution models (JSDMs) involve modelling the distribution of multiple species simultaneously, and have been growing in popularity amongst ecologists (Warton et al., 2015). The main difference between JSDMs and standard species distribution models is that the covariance between species is modelled at the same time as the response of each species to the environmental covariates (Pollock et al., 2014). This allows for information to be borrowed across species, such that the responses of rare species to the environment may be related to the responses of common species and therefore be more biologically realistic. However, the efficacy of this will depend upon the similarity of the niches of the different species (Erickson & Smith, 2023). Modelling the species variance-covariance matrix can be done in a variety of ways and some JSDM methods, i.e. those incorporating a latent variable approach, also allow for model based ordination (Hui et al., 2015).

JSDMs can, however, be high dimensional models, with each model potentially incorporating hundreds or thousands of parameters – generally constrained such that the models are identifiable. This complexity can lead to difficulties in understanding if the model fitting process has occurred successfully, and whether the model fit is truly representing the data generating process. Within this paper we present the jsdmstan package, which incorporates both JSDM data simulation and model fitting using Stan to enable greater understanding of the constraints and features of JSDMs through access to a wider suite of diagnostics and exploratory tools. Stan is a probabilistic programming language used to specify statistical models, and can be used to fit these statistical models using a variety of Bayesian algorithms including Hamiltonian Monte Carlo (Carpenter et al., 2017). The default algorithm is a dynamic Hamiltonian Monte Carlo algorithm, incorporating a series of improvements upon the original No-U-Turn Sampler (Betancourt, 2017). This provides several advantages over Gibbs samplers, adopted within other Bayesian software packages such as BUGS and JAGS, including greater efficiency and robustness in exploring the entire parameter space even in very high dimensional models. Stan also provides diagnostics that show when the entire parameter space has not been explored, and that when the model structure or fitting procedure needs to be revisited. This is particularly important in the case of JSDMs due to their high complexity and dimensionality, which can lead to poor model fits and pathological behaviour within the fitting algorithms. Therefore, the use of Stan to fit these models provides robust information for ecological inference.

## Underlying modelling approach

Within jsdmstan each JSDM is fit such that the site (*i*) by species (*j*) matrix (*M*_*ij*_) is a function of a species specific intercept (*β*_*i0*_), an environmental effect represented by species specific coefficients multiplied against the environmental covariate matrix 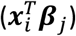 and the species covariance (*u*_*ij*_).

Optionally, an additional site-specific intercept (*α*_*i*_) can be added to adjust for total abundance or richness. This is represented in Equation 1:

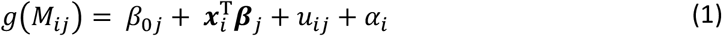

Where *g*(·)is the link function, 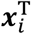is the transpose of the environmental covariate vector ***x***_*i*_, and for each taxon *jβ_0j_* is an intercept and ***β**_j_* is a vector of regression coefficients related to measured site varying predictors.

There are two ways of representing the species covariance that can be used within jsdmstan. The first is that the whole species covariance matrix can be modelled, referred to as a Multivariate Generalised Linear Mixed Model (MGLMM) throughout the jsdmstan package. Here the species covariance matrix is treated as a random draw from a multivariate normal distribution with mean **0** and covariance matrix **∑**, shown in equation 2.

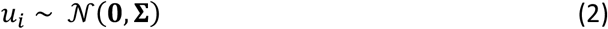

Fitting the entire covariance matrix means that the amount of time required to fit these models scales with the number of species cubed, and the data required scales with the number of species squared. This makes these models both computationally and data intensive. Alternatively, therefore, we also include Generalised Linear Latent Variable Models (GLLVMs) in which the species covariance, as included in equation 1 (*u*_*ij*_), is specified as a linear function of a set of latent variables (***z***_*i*_), equation 3. The latent variables are assumed to draw from standard normal distributions, i.e. mean 0 and standard deviation of 1. The number of latent variables is specified by the user and is generally much lower than the total number of species.

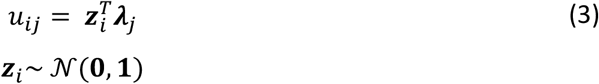

In order to make GLLVMs identifiable the upper triangle of the species loadings to latent variables (***λ**_i_*) must be fixed to zero, i.e. the first species must only load onto the first latent variable, the second species onto the first two latent variables and so on. Also, as these latent variables have the same ecological meaning whether they are positive or negative, as the sampling can lead to sign switching, which is fixed post-sampling by jsdmstan so that the signs of the latent variables do not switch within the results.

Within jsdmstan, both the GLLVM and MGLMM methods model the response of species to environmental covariates either as each response independently drawing from a prior distribution (the default), or constrained by a covariance matrix between the environmental covariates. This second option assumes that if one species is strongly positively related to multiple covariates then it is more likely that other species will either also be positively related to all these covariates, or negatively related. Mathematically this corresponds to *β*_*j*_(a *j* by *k* matrix, assuming *k* environmental covariates) being a random draw from a multivariate normal with mean 0 and covariance matrix (of dimension *k* by *k*) **Ф**.

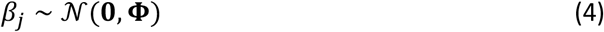

## Package description

The jsdmstan R package provides a variety of functions for both the simulation of species community data and the fitting of JSDMs to species community data. It interfaces with the rstan package for model fitting and the bayesplot package for plotting the model outputs (Gabry & Mahr, 2022; Stan Development Team, 2025). It can also interface with the loo package for leave-one-out cross validation of the model (Vehtari et al., 2020). The key functions are jsdm_sim_data which simulates data according to a specified JSDM structure and stan_jsdm which takes the data input and fits the appropriate JSDM in Stan to estimate model parameters. Both of these functions work with both a MGLMM approach and a GLLVM approach, and allow for specification of as many sites, species and environmental covariates as the user requires. The community of species can be analysed using a variety of different response families, currently including the Gaussian, Poisson, negative binomial, Bernoulli, binomial, zero-inflated Poisson and zero-inflated negative binomial families. There are a suite of functions that can be used to examine the results of each model, including summary calls and a variety of plotting functions.

### Workflow

1. **Simulation of data**. Data can be simulated using `jsdm_sim_data`, or with the aliases `mglmm_sim_data and gllvm_sim_data` which simulate data with the appropriate method. The number of species, sites, and environmental covariates can be specified by the user, in addition to directly supplying the function with environmental data to simulate the species data from. The prior distribution of the parameters used to simulate the data can also be changed through a call to `jsdm_prior()` within the simulation function.
2. **Fitting the models**. Models are fit with the `stan_jsdm` function, or alternatively the `stan_mglmm` and `stan_gllvm` aliases of this function. The Stan code used to fit each model can be accessed through supplying the relevant arguments to `jsdm_stancode()`, and the prior distributions used within the model can be modified through a call to `jsdm_prior()` within the `stan_jsdm function`.
3. **Summarising the model fits**.
  - Textual summaries. The summary and print functions return the estimated mean and quantiles of each parameter as well as the R-hat statistic and effective sample size of the bulk (Bulk-ESS) and tail (Tail-ESS) of each parameter estimate.
  - Graphical summaries. The default plot behaviour is to call `mcmc_combo` from the bayesplot package and return a traceplot and density plot for a selection of parameters. Graphical MCMC checks are provided by the `mcmc_plot` function, which interfaces to the `mcmc_ family` of functions within bayesplot. The envplot function plots the estimated environmental effects upon each species. The ordiplot function is specific to GLLVM models and plots the species or sites scores against the latent variables. The corrplot function is specific to MGLMM models and plots the modelled species correlations.
4. **Predicting from the model**. Posterior predictions can be extracted from the models using either posterior_linpred or posterior_predict, where the linpred function extracts the linear predictor for the community composition within each draw and the predict function combines this linear predictor extraction with the corresponding probability distribution to produce realisations on the original scale.
5. **Graphical posterior predictive checks**. These are supported through the pp_check function, which interfaces to the ppc_ family of functions within bayesplot. This family of functions provides a graphical way to check your posterior against the data used within the model to evaluate model fit. By default, pp_check for `jsdmStanFit` objects extracts the posterior predictions then calculates summary statistics over the rows and plots those summary statistics against the same for the original data. The summary statistic can be changed, as can whether it is calculated for every species or every site. The complementary `multi_pp_check` function plots a graphical posterior predictive check for each species.

## Examples

To demonstrate the potential usage of the jsdmstan package, we will now describe its application to two real ecological datasets. The first dataset is of records of tree genera occurrence from a survey of broadleaved woodlands where multiple plots were measured in each site. This allows demonstration of the usage of jsdmstan for binomial data where the number of trials is greater than one, and we take a MGLMM approach of estimating the entire covariance matrix. The second case study uses the widely analysed dune spider dataset, which has been used to demonstrate the usage of GLLVM models across many fitting methodologies and packages (e.g. Hui et al., 2015; Popovic et al., 2022; van der Veen et al., 2021). Here, we apply a GLLVM approach to this dataset for comparability with other methods, and discuss how jsdmstan deals with situations where there is a low amount of data for the complexity of the model.

### Broadleaved woodland data

Here we use data from a 1971 survey of broadleaved woodlands across Great Britain, which surveyed up to 16 plots per site for vegetation and soil characteristics. We model all 45 tree genera in response to soil pH, soil organic matter and three climate variables averaged over the 30 years previous to survey (annual rainfall, minimum winter temperature and maximum summer temperature). Full details on this dataset are available in Wood et al. (2015), and this data is available as part of the jsdmstan package under the name “bunce71”. The model was a MGLMM model, with all species assumed to have a binomial distribution with the number of trials equal to the number of plots per site and assuming no covariance structure within the environmental effect parameterisation (that is, the *β*_*j*_s were taken to be independent of each other). All variables were scaled to have a mean of zero and standard deviation of one before model fitting. The model was fit with default jsdmstan parameters and took approximately thirty minutes over four cores on a windows laptop. All R-hat values were under 1.01 and the number of effective samples for every parameter in both the bulk and the tail of the distribution was above 1000.

Graphical posterior checks indicate a good fit to the data, with both the number of genera per site and the number of sites per genus being recovered well by the model (Figure 1). It can be seen that most sites have between 10-15 genera present, while most genera appear in less than half the sites. The more common genera have more clearly resolved environmental responses and species covariance compared to the less common genera (Figure 2). The model can clearly find biological responses such as the preference of ash (*Fraxinus excelsior* – the only species present in the Fraxinus genus) for high pH soils and the presence of pines upon high carbon content soils (Figure 2a). The residual species correlations can be interpreted in terms of the successional gradient of the woodlands. For example, birch (Betula) is usually present early in succession while ash is present mid-to-late succession and so these species are generally negatively correlated with each other and instead found more often with genera typical of their usual successional stage (Figure 2b), e.g. birches are negatively associated with elms (Ulmus) and maples (Acer) while ash is positively associated with these (Atkinson, 1992; Thomas, 2016; Thomas et al., 2018). We cannot accurately disentangle from these results whether these are true biotic associations or represent shared response to environmental drivers, but it is likely that these represent shared response to time since disturbance or other metrics relevant to successional stage that we did not include in our model.

**Figure 1.**
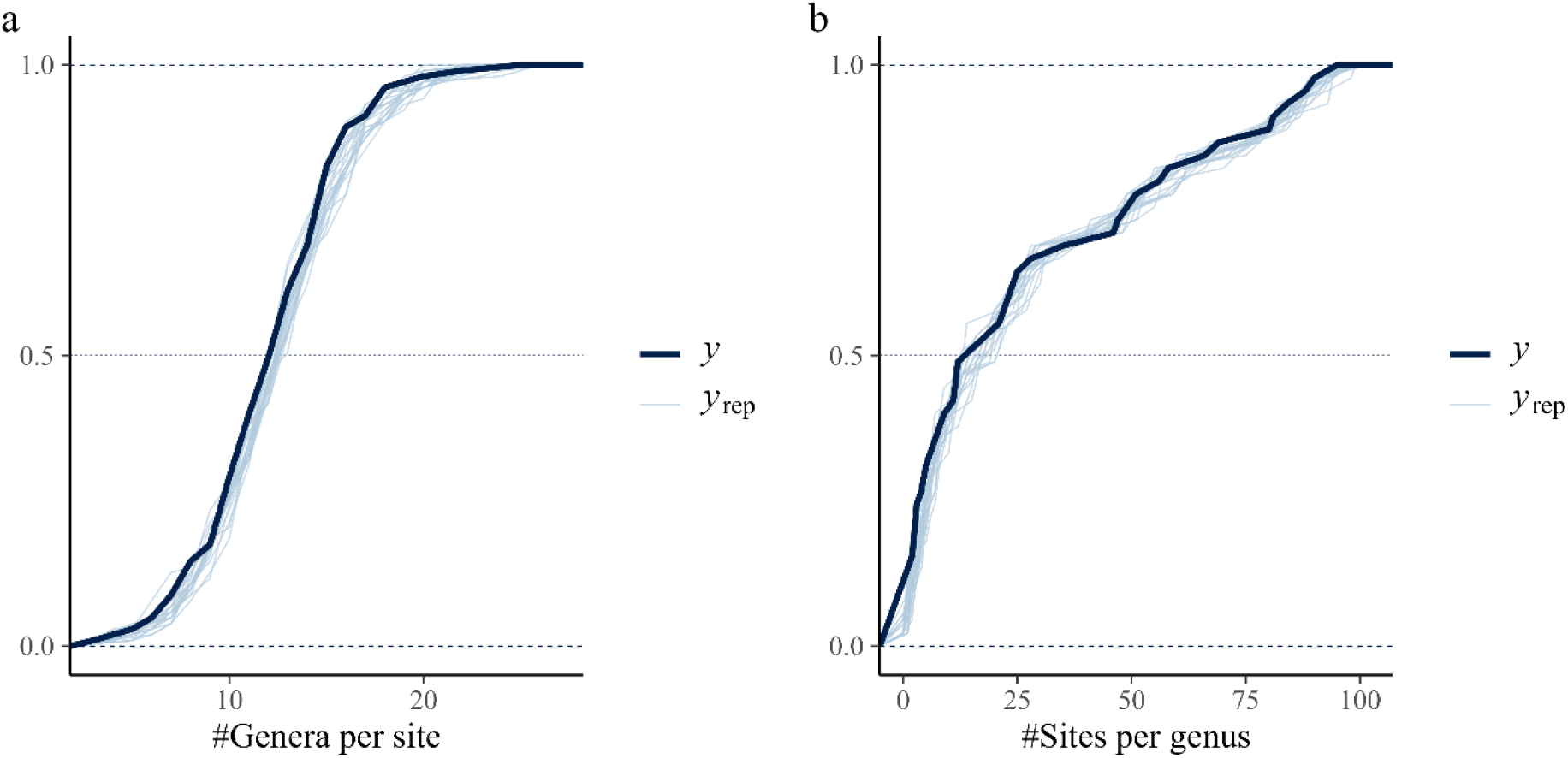
Model recovery of the number of genera per site (a) and the number of sites per genus (b). The dark blue line is the data and the light blue lines are a selection of twenty random draws from the model estimates. Close correspondence between the data and the model draws demonstrates the good fit of the model.

**Figure 2.**
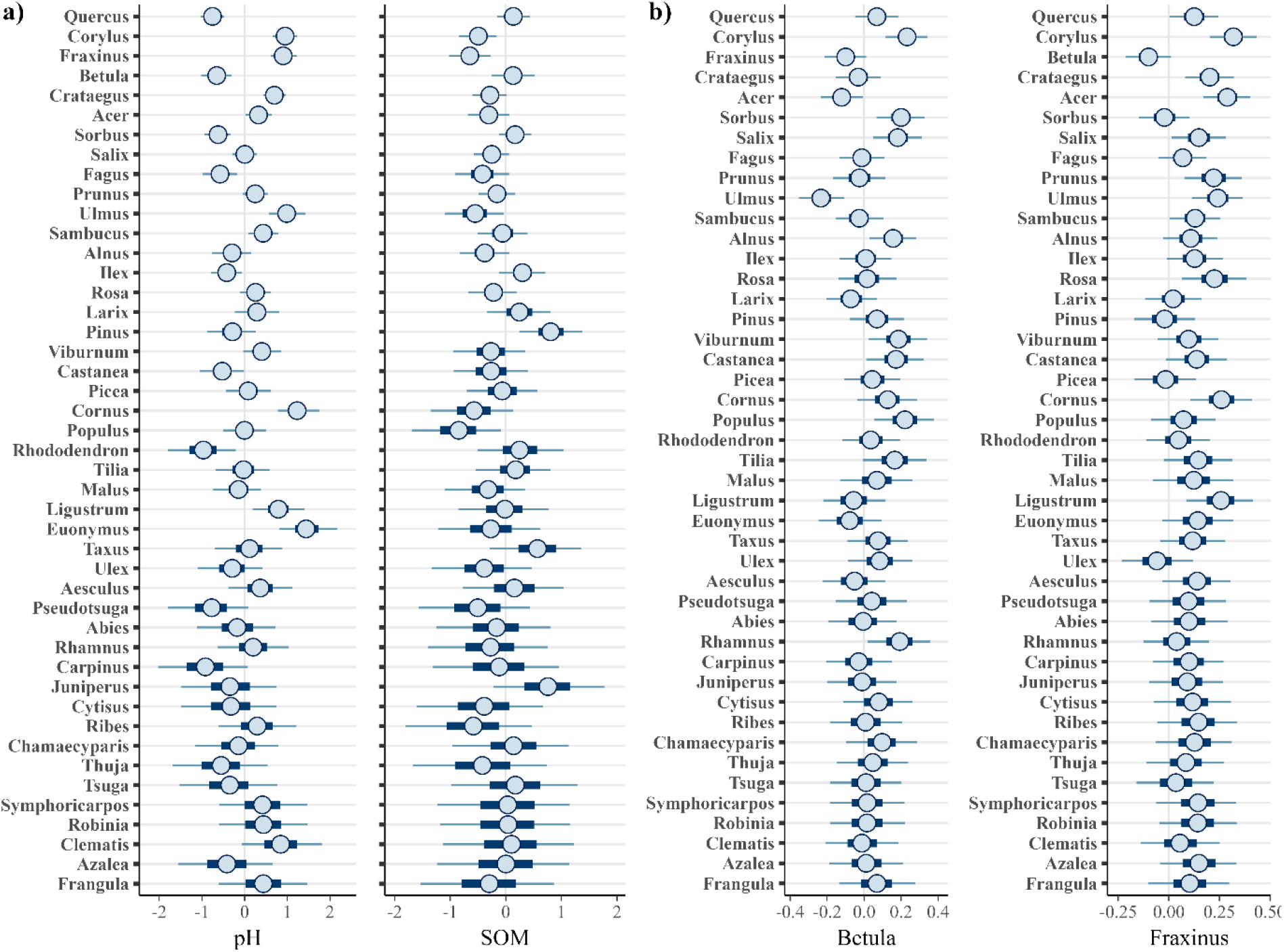
The estimated effects of soil pH, and SOM upon each genus (a) and the estimated correlations between genera for Betula and Fraxinus (b). Genera are listed in order of frequency of appearance, with the genera that appear in the most sites at the top of the figure. The estimates are shown as the median (circle), 50% interval (thick blue line) and the 90% interval (thin blue line).

### Dune spider data

Here we use data from a pitfall trap survey of spiders in Dutch dunes to demonstrate fitting a GLLVM. This data consists of 12 spider species measured over 28 dune sites with 6 environmental covariates measured at the same sites and is available within the mvabund R package (van der Aart & Smeenk-Enserink, 1975; Wang et al., 2012). The model was run with two latent variables (from equation 3, ***z***={*z*_1_*z*_2_}), a negative binomial response and default jsdmstan parameters. This took approximately 2 minutes on a windows laptop over four cores, with all parameters having R-hat less than 1.01 and effective sample sizes of the bulk and tail being over 1000.

Initial attempts to run this model with all six environmental covariates showed that the loadings onto the latent variables for both the species and the sites trended towards a diagonal with high uncertainty around individual estimates (Figure 3a-b). This indicates that the model was unable to fully resolve the latent variables, likely due to there not being enough information in the data to separate out the latent variable effects from the environmental covariate effects. Therefore, the model was run again with two environmental covariates (soil.dry and herb.layer) and two latent variables. Again, this model had all parameters with R-hat less than 1.01 and effective sample sizes of the bulk and tail being over 1000. This model resolved the latent variable effects, with no diagonal loadings apparent (Figure 3c-d). It is worth noting that both models successfully recovered the distribution of abundance and richness across the sites (Figure 4), indicating that fit to data alone is insufficient to detect all potential model fit issues.

**Figure 3.**
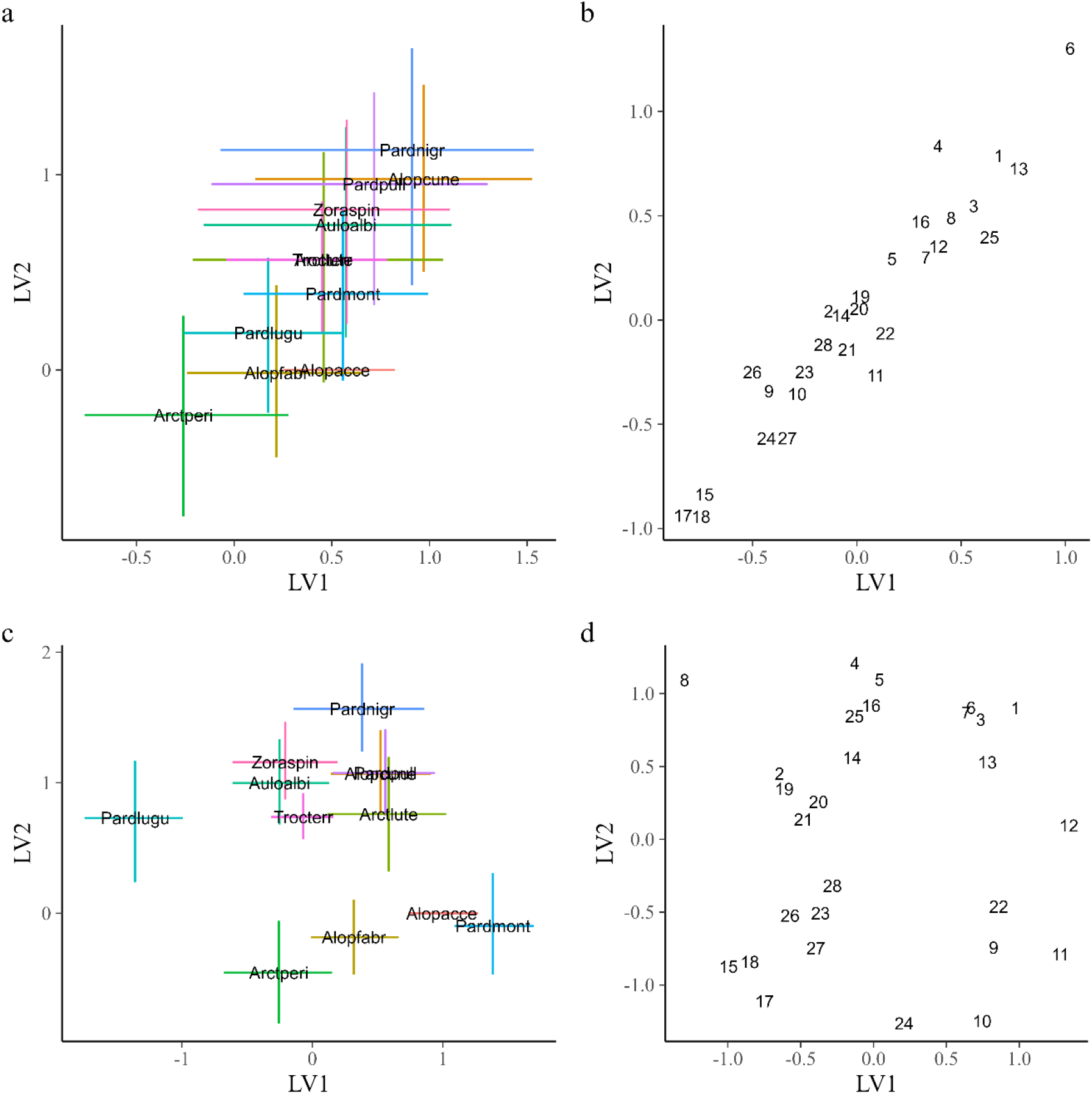
The species (a, c) and site (b, d) mean loadings onto the two latent variables for the model with six environmental covariates (a, b) and the model with two environmental covariates (c, d). For species only the 50% range of the loadings are also shown as lines upon the graph.

**Figure 4.**
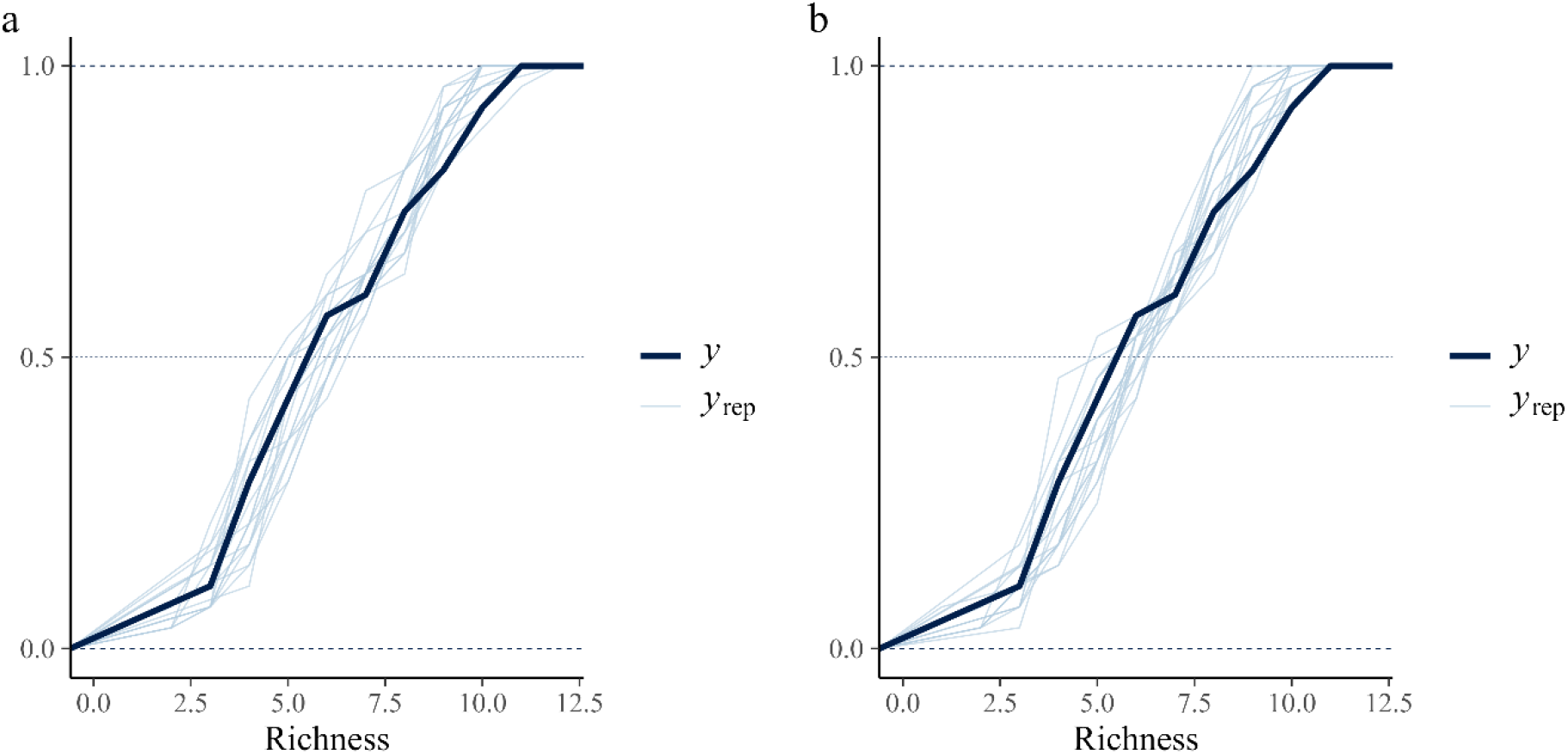
Model recovery of species richness for the model with all environmental covariates (a) and the model with two environmental covariates (b). The dark blue line is the data and the light blue lines are draws from the model estimate.

Ecological interpretation of the second model suggests that the separation out of pardmont, arctperi, alopfabr, alopacce within this second model could be due to a shared positive response to moss. As moss is no longer included as an environmental predictor, the latent variables are free to take up the explanatory power of the moss gradient, which is consistent with the understanding of latent variables generally being measures of missing environmental variables (Vallé et al., 2024). Balancing the ability to include multiple environmental predictors and more general latent factors in low data contexts is a challenge, and the most appropriate choice will depend on the intended use of any particular model.

## Conclusions

The jsdmstan package offers a new way to fit joint species distribution models, through using the Stan probabilistic programming language with advanced model diagnostics and graphical checks incorporated. It supports multiple ways of specifying the species interactions, including both specification of the full covariance matrix and specification of the covariance matrix through a latent variable model. Future directions for this package include incorporating spatio-temporal and other random effect information into the model, as well as working on speed improvements to model fitting and comparison. We expect that the jsdmstan package will provide ecologists with the tools to fit a variety of joint species distribution models; therefore achieving greater understanding of both how well these models are describing the data and how environmental effects and species associations are affecting species communities.

## Acknowledgements

This work would not have been possible without the Stan forums community, particularly Jim Downie who developed Stan code for a GLLVM. Thanks are also due to Adam Kimberley and Simon Smart who offered advice for understanding the woodland data and model results, and to Pete Henrys for his feedback on the modelling approach. This research was supported by NERC, through the UKCEH National Capability for UK Challenges Programme NE/Y006208/1.

## Data and code availability

The Bunce woodland data is available from the Environmental Information Data Centre under (Kirby et al., 2013c, 2013a, 2013b). The climatic data is available to download from the NERC EDS Centre for Environmental Data Analysis (Met Office et al., 2022). The Bunce woodland data used in this paper are available within the jsdmstan package presented here, and the spider data is available within the mvabund R package (Wang et al., 2012). The jsdmstan R package is available at https://github.com/NERC-CEH/jsdmstan (Seaton, 2025), and the R code for the case studies included in the Supplementary Material below and at https://github.com/NERC-CEH/ds-toolbox-notebook-jsdmstan.

## Supplementary Material: Code for Case Studies

~~~
library(jsdmstan)
library(dplyr)
library(ggplot2)
library(patchwork)
set.seed(3757892)
## ----read in bunce71 data
data(“bunce71”)
## ----scale b71 data
bunce71_env <-mutate(bunce71$env, across(pH:WinterMinTemp, scale))
bunce71_abund <-bunce71$abund[,names(sort(colSums(bunce71$abund>0),
                                                        decreasing = TRUE))]
## ----bw model fit, cache = TRUE
bwmod <-stan_mglmm(∼Rainfall + SummerMaxTemp + WinterMinTemp + pH + SOM,
          data = bunce71_env, Y = bunce71_abund,
          Ntrials = bunce71_env$Nplots,
          family = “binomial”,
          cores = 4)
bwmod
## ----bw pp_check
pp_check(bwmod)
p1 <-pp_check(bwmod, plotfun = “ecdf_overlay”,
       summary_stat = function(x) sum(x>0), ndraws = 20) +
     labs(x = “#Genera per site”)
p2 <-pp_check(bwmod, plotfun = “ecdf_overlay”, calc_over = “species”,
       summary_stat = function(x) sum(x>0), ndraws = 20) +
    labs(x = “#Sites per genus”)
p1+p2 + plot_annotation(tag_levels = “a”)
## ----bw multi_pp_check
multi_pp_check(bwmod, plotfun = “ecdf_overlay”, species = 1:9)
multi_pp_check(bwmod, plotfun = “ecdf_overlay”, species = 10:18)
multi_pp_check(bwmod, plotfun = “ecdf_overlay”, species = 19:27)
multi_pp_check(bwmod, plotfun = “ecdf_overlay”, species = 28:36)
multi_pp_check(bwmod, plotfun = “ecdf_overlay”, species = 37:45)
## ----bw envplot
envplot(bwmod, widths = c(1.5,1,1))
## ----bw corrs
corrplot(bwmod, species = c(“Betula”,”Fraxinus”,”Quercus”,”Corylus”))
e1 <-envplot(bwmod, widths = c(1.4,1), preds = c(“pH”,”SOM”))
c1 <-corrplot(bwmod, species = c(“Betula”,”Fraxinus”))
ggpubr::ggarrange(e1,c1, labels = c(“a)”,”b)”,”c)”,”d)”))
## ----get spider data
data(“spider”, package = “mvabund”)
spider$x <-scale(spider$x)
## ----run spider mod
spmod <-stan_gllvm(X = spider$x, Y = spider$abund, D = 2,
                           family = “neg”, cores = 4)
spmod
## ----sp pp_check
p1 <-pp_check(spmod, plotfun = “ecdf_overlay”, ndraws = 20) +
    labs(x = “Abundance”) +
    coord_cartesian(xlim = c(0,750))
p2 <-pp_check(spmod, plotfun = “ecdf_overlay”,
           summary_stat = function(x) sum(x>0), ndraws = 20) +
    labs(x = “Richness”)+
     coord_cartesian(xlim = c(0,12))
p1 + p2 + plot_annotation(tag_levels = “a”)
## ----sp multi_pp_check
multi_pp_check(spmod, plotfun = “ecdf_overlay”)
## ----sp envplot
envplot(spmod, widths = c(1.2,1,1))
## ----sp ordiplot
o1 <-ordiplot(spmod, ndraws = 0, geom = “text”, size = c(3,1),
                     errorbar_linewidth = 0.5, errorbar_range = 0.5) +
    theme(legend.position = “none”)
o2 <-ordiplot(spmod, ndraws = 0, geom = “text”, type = “sites”, size =
c(3,1),
                     errorbar_range = NULL)
o1 + o2 + plot_annotation(tag_levels = “a”)
## ----update sp mod with fewer covariates
spmod2 <-update(spmod, newX = spider$x[,c(“soil.dry”,”herb.layer”)],
                       cores = 4)
spmod2
## ----sp2 pp_check
p1b <-pp_check(spmod2, plotfun = “ecdf_overlay”, ndraws = 20) +
    labs(x = “Abundance”) +
    coord_cartesian(xlim = c(0,750))
p2b <-pp_check(spmod2, plotfun = “ecdf_overlay”,
        summary_stat = function(x) sum(x>0), ndraws = 20) +
    labs(x = “Richness”)+
    coord_cartesian(xlim = c(0,12))
p1b + p2b + plot_annotation(tag_levels = “a”)
p2 + p2b + plot_annotation(tag_levels = “a”)
## ----sp2 multi_pp_check
multi_pp_check(spmod2, plotfun = “ecdf_overlay”)
## ----sp2 envplot
envplot(spmod2, widths = c(1.2,1))
## ----sp2 ordiplot
o1v2 <-ordiplot(spmod2, ndraws = 0, geom = “text”, size = c(3,1),
                       errorbar_linewidth = 0.5, errorbar_range = 0.5) +
    theme(legend.position = “none”)
o2v2 <-ordiplot(spmod2, ndraws = 0, geom = “text”, type = “sites”,
                     size = c(3,1), errorbar_range = NULL)
## ----sp all ordiplots
o1 + o2 + o1v2 + o2v2 + plot_annotation(tag_levels = “a”)
~~~

